# *In-situ* Real-time Field Imaging and Monitoring of Leaf Stomata by High-resolution Portable Microscope

**DOI:** 10.1101/677450

**Authors:** Prashant Purwar, Junghoon Lee

## Abstract

Stomata, functionally specialized micrometer-sized pores on the epidermis of leaves (mainly on the lower epidermis), control the flow of gases and water between the interior of the plant and atmosphere. Real-time monitoring of stomatal dynamics can be used for predicting the plant hydraulics, photosensitivity, and gas exchanges effectively. To date, several techniques offer the direct or indirect measurement of stomatal dynamics, yet none offer real-time, long-term persistent measurement of multiple stomal apertures simultaneously of an intact leaf in a field under natural conditions. Here, we report a high-resolution portable microscope-based technique for *in situ* real-time field imaging and monitoring of stomata. Our technique is capable of analyzing and quantifying the multiple lower epidermis stomal pore dynamics simultaneously and does not require any physical or chemical manipulation of a leaf. An upward facing objective lens in our portable microscope allows the imaging of lower epidermis stomatal opening of a leaf while upper epidermis being exposed to the natural environment. Small depth of field (~ 1.3 μm) of a high-magnifying objection lens assists in focusing the stomatal plane in highly non-planar tomato leaf having a high density of trichome (hair-like structures). For long-term monitoring, the leaf is fixed mechanically by a novel designed leaf holder providing freedom to expose the upper epidermis to the sunlight and lower epidermis to the wind simultaneously. In our study, a direct relation between the stomatal opening and the intensity of sunlight illuminating on the upper epidermis has been observed in real-time. In addition, real-time porosity of leaf (ratio between the areas of stomatal opening to the area of a leaf) and stomatal aspect ratio (ratio between the major axis and minor axis of stomatal opening) along with stomatal density have been quantified.

## INTRODUCTION

Evolution and natural growth of plant life are based on the continuous spectrum of the sunlight [1–3]. Major plant growth processes such as transpiration, photosynthesis and respiration require a controlled gaseous exchange between the interior of the plant and atmosphere. Stomata, functional pores found at the epidermis of the leaf, regulate this gaseous exchange during growth processes by controlling the aperture of stomatal opening [4]. Stomatal opening is predominantly affected by sunlight intensity (SLI) falling on the upper epidermis of a leaf [3, 5]. Changes in stomatal aperture are determined by two major approaches: directly by measurement of the stomatal apertures and indirectly by measurement of leaf transpiration. To date, several techniques offer the direct or indirect measurement of stomatal aperture or opening/closing information of stomatal pores, yet none offer real-time, long-term persistent measurement of multiple stomal apertures simultaneously of an intact leaf under natural conditions.

Porometer, an instrument to measure the stomatal resistance (or conductance) against the gaseous exchange only provides the mean pore size of stomata (if stomatal density is known) and not suitable for long-term persistent stomatal monitoring [6]. Plasmolysis and electro-mechanical sensor techniques indicates only the open or closed status of stomata [7, 8]. Infrared thermography based technique are limited to the bulk analysis of stomata behavior and suffers for several limitations such as accuracy, thermal drift of imager, unreliable references for thermal indices etc. [9–12]. On the other hand, direct measurement mainly utilizes microscopic images of a detached leaf or imprints of an intact leaf, which often requires invasive manipulations (physical or chemical) of a leaf [13–15]. Optical microscopy of abaxial stomata of an intact leaf by a conventional reflected microscope requires 180 degrees folding of a leaf to face the abaxial side towards the objective lens. Such folding often damages the leaf and restrict the upper epidermis exposing to the light source [8]. Mold impression technique, which utilizes the permanent impression of stomata on silicon rubber, does not provide the accurate measurement of stomata [16–19]. Fluorometric measurement of individual stomatal activity utilizes water-responsive UV treated polymer coating on the abaxial surface of the leaf. However, polymerization of a water-responsive polymer may damage the leaf. In addition, quantification of stomatal opening requires expensive optical and fluorescence microscopes[20–22].

Herein, we developed a portable high-resolution microscopy-based technique for *in situ* real-time field imaging and monitoring of stomata at a single stoma level analysis. Our technique is capable of analyzing and quantifying the multiple stomal pore dynamics simultaneously and does not require any manipulation of a leaf. A novel leaf holder has been designed and fabricated to hold the leaf for long-term monitoring of stomata. Leaf holder allows exposing the upper epidermis to the sunlight and lower epidermis to the wind simultaneously. After analyzing the dynamics of stomatal aperture in tomato plant leaves, it was found that the aperture of a stomatal pore is a direct function of the intensity of sunlight illuminating on the upper epidermis. In addition, stomatal density, porous area of the leaf (ratio between the areas of stomatal opening to the area of the leaf) and stomatal aspect ratio (ratio between the major axis and minor axis of stomatal opening) were also quantified.

## 2. MATERIAL AND METHODS

### 2.1 Experimental Setup

A high-resolution portable reflected microscope having an upward facing objective lens and vertical illumination is shown in Fig. 1. Our microscope consists of a commercial 40× magnification finite microscope objective lens (NA 0.6), an image sensor (1600 × 1200 pixels, pixel size 3 μm × 3 μm, Eyecam), a plate type beam splitter (BS, 25 mm in diameter, 30R/70T, Edmund Optics) and light-emitting diode (LED) based light source. The light source consists of a two-watt surface-mounted white LED and an aspherical condenser lens (Ø: 25 mm, focal length: 20 mm, NA 0.6). The image sensor was placed close to the objective lens for compactness of the setup (Fig. 1A and 1B) and tube lens was not used. Microscopic components were optically aligned with the help of 3D printed structures (Fig. 1C). To hold the leaf surface parallel to the microscope-imaging plane, a leaf holder has been designed and fabricated.

**Fig. 1.**
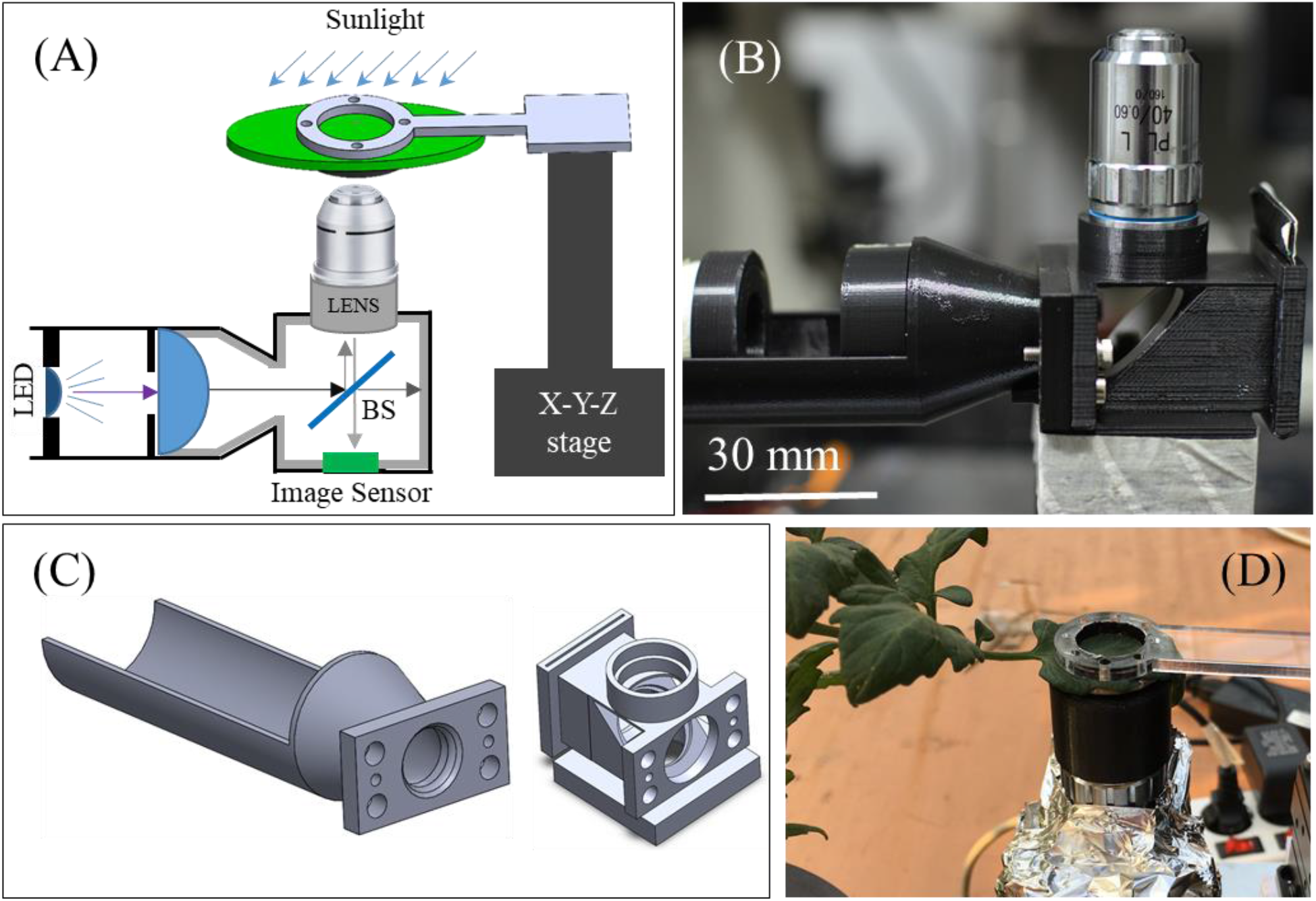
Experimental setup (A) Schematic of the portable reflected type microscope having upward facing objective lens and novel leaf holder. Our setup is capable of direct real-time imaging and monitoring of lower epidermis stomata in field (B) Customized compact reflected microscope consists of a commercial finite 40× objective lens, BS and LED-based light source in 3D printed support frame (light source holder and lens holder). (C) Computer-aided design (CAD) of light source holder and lens holder. All the cavities of lens holder were lined with black color velvet paper from inside to reduce the reflection of light from walls of cavities. The whole microscope was covered by aluminum foil to ensure that ambient light should not interfere with the microscopic optical system in sunlight (D) Direct real-time imaging of lower epidermis stomata in field.

### 2.2 Leaf Holder

A novel leaf holder has been fabricated with the help of a laser cutter (Universal Laser System USA) and 3D printer (DP200, Sindoh Korea) as shown in Fig 2. Leaf holder consists of four parts, named as upper leaf holder, lower leaf holder, leaf cap and a lens cap (Fig. 2A and 2B). Upper leaf holder is made of 3 mm thick biocompatible polymethylmethacrylate (acryl) sheet and allows a 15 mm diameter area of the leaf for microscopy [23, 24]. The upper leaf holder is attached to an x-y-z stage to focus the leaf surface for microscopic imaging. A lower leaf holder of thickness 1.5 mm is fabricated with a black color biocompatible polylactic acid (PLA) material to restrict the ambient light reaching to the sample from backside [25, 26]. The thickness of the lower leaf holder was determined after considering the distance between the leaf surface and an objective lens (~2.5 mm) when the lower epidermis was in focus. To avoid the interference of ambient light reflected from lens surface with the optical system of the microscope, the lens was also covered with a black color lens cap made of PLA material. V-shaped notches (2 mm wide) were patterned on lower leaf holder to pass the wind & moisture and restrict the sunlight across the abaxial leaf area under the investigation. The sample (leaf) was anchored in between the upper and lower leaf holders with the help of four small permanents neodymium magnets embedded in the upper (Ø 2 mm, thickness 1.5 mm) and lower (Ø 3 mm, thickness 1.5 mm) leaf holders. During microscopy, upper leaf surface was closed by leaf cap to restrict the interference of ambient sunlight with the optical system of the microscope as shown in Fig. 2C and Fig. 2C’.

**Fig. 2.**
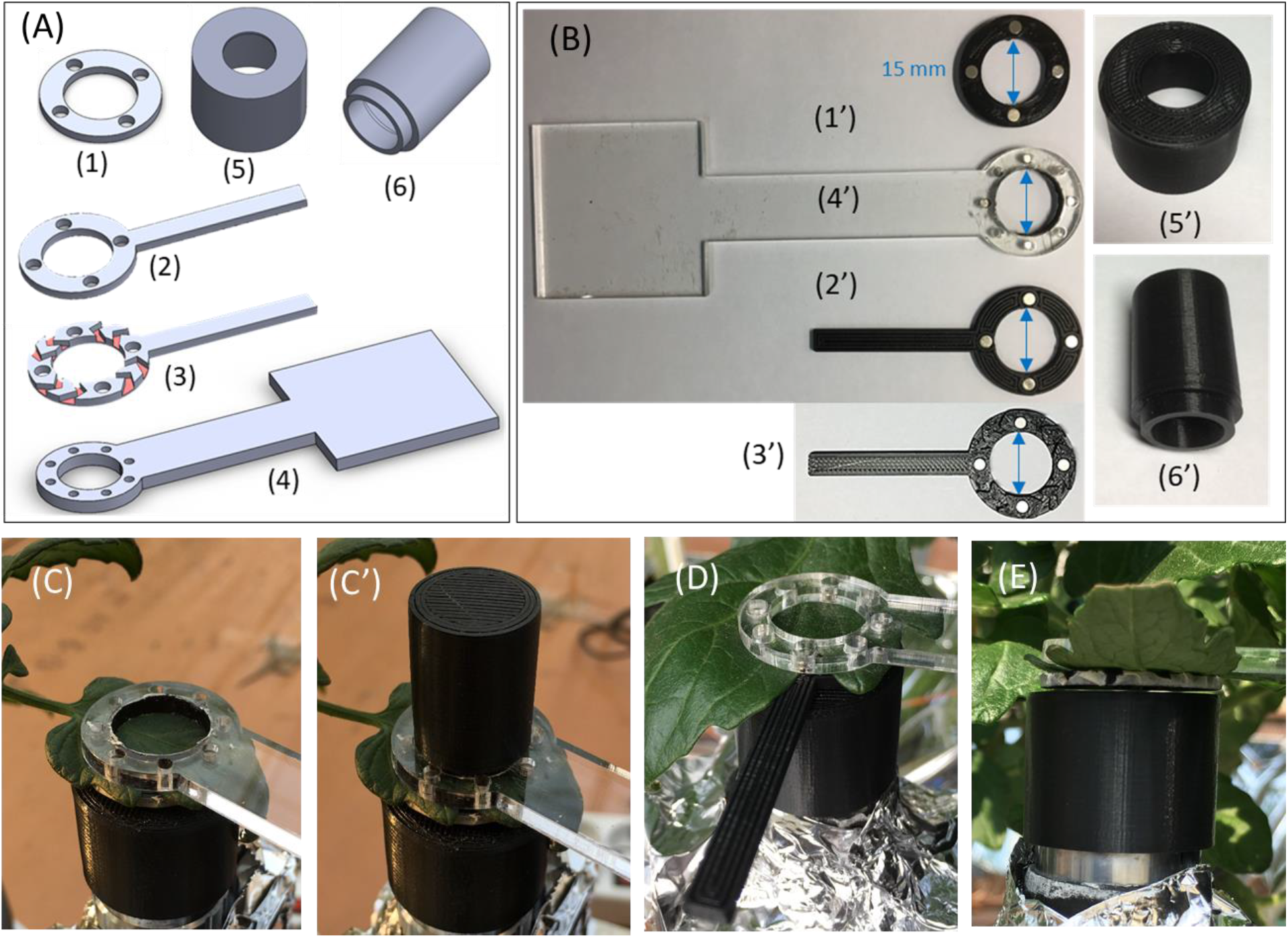
Leaf holder: a device to hold and stabilize the leaf for long-term stomata imaging. (A) CAD design of leaf holder components. Upper leaf holder (4) is made of 3 mm thick transparent acryl sheet to allow continuous monitoring of stomata while the upper surface of the leaf exposed to the sunlight. However, 1.5 mm thick lower leaf holder (1, 2 or 3) was fabricated with a 3D printer using black color PLA material with and without a handle. V-shaped notches of width 2 mm have been designed to allow the wind blow and restrict the ambient light at the abaxial layer of the leaf. Small permanent neodymium magnets embedded in the upper and lower leaf holder helps to hold the leaf even if during the wind blow. (B) 3D-printed structures of leaf holder (C) Leaf holder in normal condition (C’) Use of lens cap to avoid ambient sunlight during microscopy. (D) Lower leaf holder with handle. Handle helps to attach the lower leaf holder to the upper part without damaging the biological structures of the lower epidermis of the leaf. (E) A gap of 0.5~1 mm between the lens cap and lower leaf holder to move the sample freely for stomata imaging.

### 2.3 Tomato Plant Cultivation

Stomata imaging of tomato leaf was conducted in a transparent vinyl film covered greenhouse under natural environment. Experiments were conducted on two ways grew tomato plants, one having controlled irrigation of nutrients based on solar radiation and another having the irrigation of nutrients on a daily basis. Tomato plants were planted in Rockwool cubes (10 × 10 × 6.5 cm, UR Rockwool, Korea) and sufficiently saturated with the standard nutrient solution for tomato plants (Glasshouse Crops Research and Experiment Station at Naaldwijk, Netherlands). Nutrient solution was supplied with the help of a proportional control method of integrated solar radiation using a drip irrigation system (Netajet, Netafim Korea, Korea). Experiments were conducted on six weeks old tomato plants. During the experiment, wind velocity was negligible.

### 2.4 Electron Microscopy and Surface Profile Measurement

A field-emission scanning electron microscope (FESEM, Carl Zeiss SUPRA Germany) and surface profiler (Nanofocus μsurf) was used to observe the leaf surface texture and measure the surface roughness of the leaf respectively. To prepare the leaf sample for FESEM imaging, a fresh detached leaf was cut into the 2 mm × 2 mm pieces and submerged immediately into a Karnovsky fixative media for 2 hours at room temperature. After primary fixation, leaf pieces were washed thoroughly in 0.05M sodium cacodylate buffer 3 times followed by in distilled water 2 times. To dehydrate the sample, the sample was washed in graded ethanol (30%, 50%, 70%, 80%, 90% and 100% (3 times)) each for 10 min. Finally, the leaf samples were dried in Critical Point Dryer (EM CPD300, LEICA) before electron microscopy.

### 2.5 Fabrication of Rectangular Grid Pattern

Rectangular grid patterns of a negative photoresist, SU-8 3005 (Microchem USA) was fabricated on a 4-inch silicon wafer by a single-mask photolithography process. First, photoresist SU-8 was spin-coated on a clean silicon wafer at 500 RPM and 4000 RPM speed for 30 seconds and 60 seconds respectively, followed by a soft bake process at 65°C for 180 sec. Next, the wafer was exposed to 150 mJ/cm^2^ UV (365 nm) source followed by a post-exposure bake at 95°C for 60 seconds. Further, the sample was immersed in MicroChem’s SU-8 developer for 120 seconds to develop the photoresist and rinsed with deionized water thoroughly. Finally, the sample was hard bake at 150° for 60 seconds for good adhesion of developed SU-8 grid pattern on a silicon wafer.

### 2.6 Measurement of Sunlight Intensity

The intensity of sunlight was measured with the help of a light-meter (Lutron LX-1108 Taiwan) in unit ‘Lux’. To measure the maximum SLI inside the greenhouse, the sensor of the light-meter was faced towards the sun and the maximum value was recorded. On the other hand, SLI value on the leaf surface was estimated by placing the sensor of light-meter close and parallel to the leaf surface.

### 2.7 Measurement of Optical Resolution

An optical resolution of the portable microscope was measured in term of half-pitch spatial resolution (HSR) by an extreme microscope resolution target, 1951 USAF (Amazon.com). Microscope resolution target has the smallest HSR value of 0.137 μm. Resolution target contains 100 nm thick three equidistant vertical and horizontal direction chrome bars of different HSR values increasing in width by ~12.4 % of the previous value (HSR value: 0.137 μm to 31.250 μm) plated on quartz background. An optical resolution of a microscope (with a 40× objective lens, NA 0.6) was determined after distinguishing the three vertical and/or horizontal parallel bars of minimum HSR value.

### 2.8 Quantification of Stomatal Density

Microscopic images of stomata were captured by a portable microscope and analyzed in ‘ImageJ’ software. Stomatal density (stomatal number per unit leaf area) was calculated by averaging the number of stomata observed at five different parts of a leaf (lower epidermis). To distinguish the stomatal complex from other epidermis structures, stomatal counting was performed when stomata were widely open.

## 3. RESULTS AND DISCUSSION

### 3.1 Compact Optical Microscope Imaging and Optical Resolution

Microscale imaging of rectangular grid patterns (50 μm × 50 μm × 5 μm) of photoresist SU-8 on a silicon wafer and measurement of an optical resolution of our microscope is shown in Fig. 3. The image quality taken by our microscope was compared with the image of a Nikon eclipse reflected microscope (20× objective lens, NA 0.45) using of rectangular grid patterns. Image quality obtained by both microscopes were comparable and no distortion has been observed in the image obtained by our microscope. However, FOV (350 μm × 262 μm) of our microscope was smaller than the FOV (670 μm × 500 μm) provided by a Nikon microscope (20× objective lens, NA 0.45). Still, the imaging area of 350 μm × 262 μm was enough to monitor 20-30 stomata simultaneously in the abaxial layer of the tomato leaf. For compactness of the microscope, the image sensor was placed close to an objective lens without using the tube lens between an objective lens and image sensor. This further reduces the optical attenuation in the image formation owing to the optical systems of the tube lens, which helps to obtain a sharp image of an object. Without tube lens, the image of an object should be distorted in large FOV, however, in small FOV (350 μm × 262 μm) of our microscope, no distortion has been observed in the image of rectangular grid patterns as shown in Fig. 3A.

**Fig. 3.**
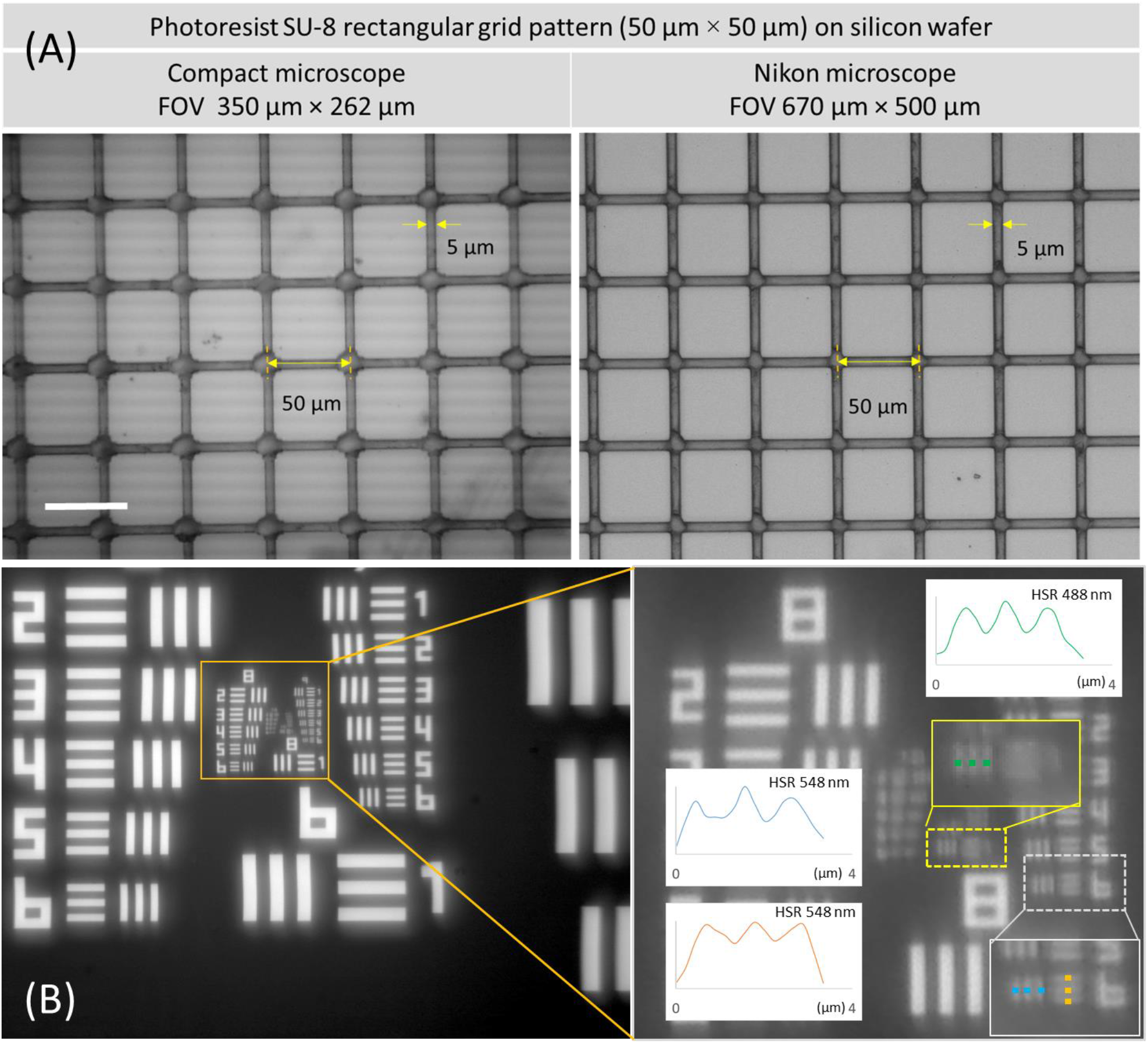
Microscale imaging and optical resolution (A) Imaging of microscale rectangular grid patterns (50 μm × 20 μm × 5 μm) of photoresist SU-8 on a silicon wafer. Image quality obtained by compact microscopic setup is comparable to the standard Nikon reflected microscope. Band noise (horizontal lines) in the image obtained by our microscope was due to overexposure of image sensor. Scale bar 50 μm (B). Optical resolution in term of HSR value. HSR value was 0.548 μm (FOV 350 μm × 262 μm) when vertical and horizontal both direction lines were clearly distinguishable and 0.488 μm when one-direction lines were distinguishable. We believe that the background vibrations, smaller distance lens and image sensor were the reasons not clearly visualize both direction lines at HSR value of 0.488 nm.

The optical resolution of the portable microscope was measured in terms of half-pitch spatial resolution (HSR) by extreme microscope resolution target, 1951 USAF. At HSR value of 0.548 μm (FOV 350 μm × 262 μm), both direction lines (vertical and horizontal) were clearly distinguishable. However, a lower HSR value (0.488 μm) was obtained when only one-direction lines were distinguishable. We believe that the background vibrations was one of the reason not to clearly visualize both direction lines at HSR value of 488 nm (Fig. 3). Overexposing the image sensor by a LED-based vertical light source produced the band noise (horizontal lines) in the image obtained by our microscope (Fig. 3A).

### 3.2 Tomato Leaf Surface Profile

Understanding the surface texture of a leaf is important for the optical microscopy of stomata. Surface profile measurement and high-resolution imaging of tomato leaf were conducted by 3D surface profiler (at 176× optical magnification) and FESEM (at 500× magnification) respectively (Fig. 4). Results showed a highly non-planar surface of tomato leaf with an height profile of 172 μm in FOV of 800 μm × 800 μm. In addition, FESEM images revealed a dense population of trichome, a hair and granular like structures on the surface of a leaf. Non-planar surface of the leaf limits focusing the stomata in large FOV. Hence, multiple images focusing at the different stomatal plane in given FOV were captured to analyze the stomatal opening and stomatal density. Moreover, the presence of high-density trichome on leaf surface further hinders in obtaining the high contrast images of stomata. However, the use of a 40× microscopic objective lens having a small depth of field (~ 1.3 μm) helps to focus on the stomatal plane even if in the presence of high-density trichome. Tomato stomata imaging in FOV of 350 μm × 262 μm is shown in Fig. 5D and 5D’.

**Fig. 4.**
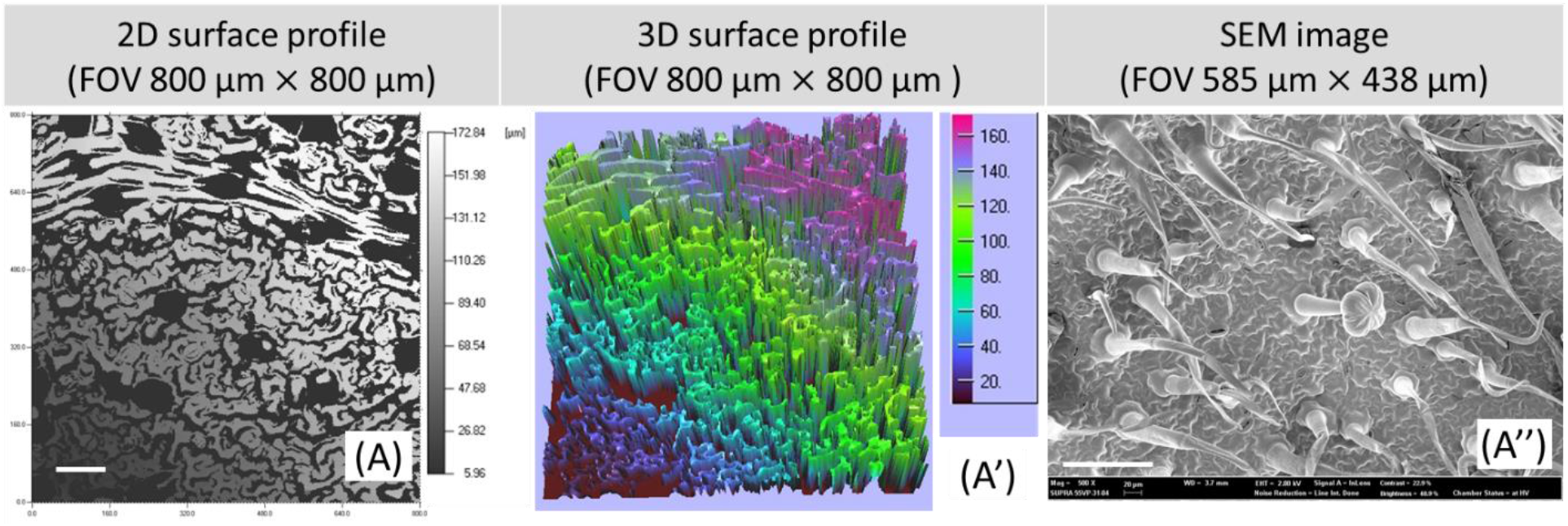
Surface profile measurement and electron microscopy tomato leaf. (A and A’) 2D and 3D surface profile measurement by 3D surface profiler of tomato leaf at 176× optical magnification. A height profile map demonstrates higher non-planarity of tomato leaf of 172 μm (FOV of 800 μm × 800 μm). Scale bar: 100 μm. (A’’) FESEM image of the lower epidermis of a tomato leaf at 500× magnification showing highly dense population of trichome. Trichome are hair like structures, which makes the surface further difficult for imaging. Scale bar: 100 μm

**Fig. 5.**
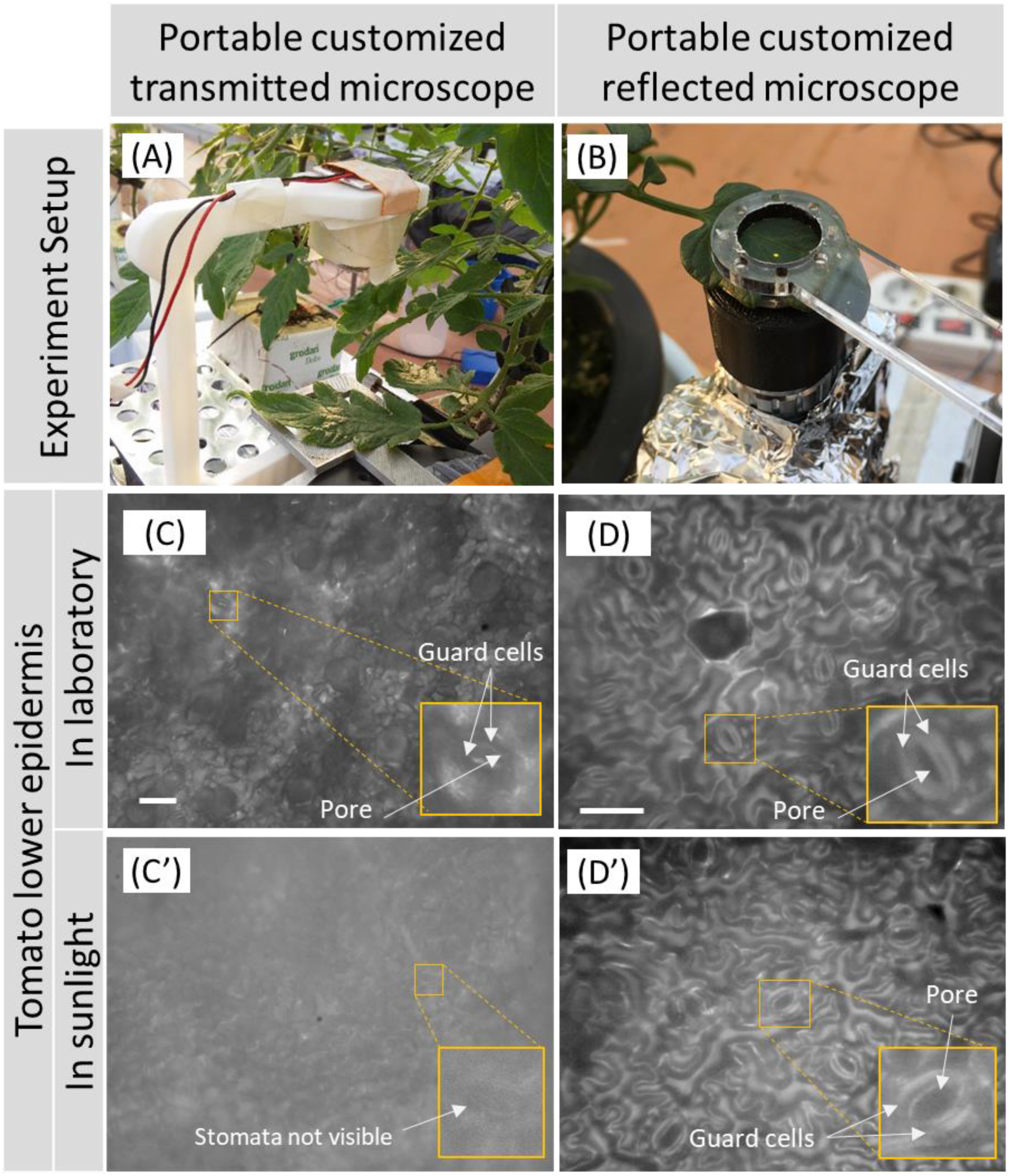
Stomata imaging by compact transmitted and reflected microscopes in laboratory and field environment. Experimental setup for stomata imaging by (A) a portable transmitted type microscope (B) a portable reflected type microscope. Observation of tomato stomata of lower epidermis when imaged in the laboratory environment by (C) transmitted microscope. Scale bar: 100 μm (D) reflected microscope. Scale bar: 50 μm and in the field by (C’) transmitted microscope (D’) reflected microscope. In case of a transmitted microscope, stomata observation was only possible in a laboratory environment since ambient light in field interfere with the optical system of a microscope. The image quality of stomata imaging was comparable when observed by reflected microscope in a laboratory as well as field environment. Stomata and guard cells, both, were clearly visible.

### 3.3 A Need of Reflected Microscope with Vertical Illumination for *in situ* Stomata Imaging

An interaction of ambient light with the optical system of a microscope may result in poor imaging of an object. A comparative study of lower epidermis stomata imaging of a tomato leaf (leaf thickness 150 μm-250 μm, excluding leaf veins) by a portable transmitted and reflected microscopes in dark (inside laboratory) and bright (inside the greenhouse during daytime) environment is shown in Fig. 5. A reflected microscope provides high-quality images of abaxial stomata in the laboratory as well as in the field environment. Both microscopes consist of 40× finite objective lens; an image sensor and white LED-based imaging source. Details of a portable transmitted microscope are mentioned somewhere else [27]. In case of the transmitted microscopy, imaging of the lower epidermis without any interaction between ambient light and a microscopic optical system is difficult while exposing the upper epidermis to the ambient light in the field. Moreover, the use of a high-intensity light source to counter the effect of ambient light may damage the leaf and affect the working of stomata. In addition, imaging of lower epidermis stomata by a transmitted microscope should also have footprints of several upper and intermediate layers of the leaf, which will produce a poor contrast image of lower epidermis stomata (Fig. 5C). As a result, a transmitted microscope is not suitable for stomata imaging. Hence, a reflected microscope capable of lower surface imaging of a leaf should be used for stomata imaging.

Unlike commercial reflected microscopes, our portable microscope consists of an upward facing objective lens and microscope can be placed under the leaf surface easily. Use of side illumination (specular and diffuse illumination) can produce shadows on the rough surface of a leaf, a vertical Kohler illumination source was used to obtain a clear and contrast image of stomatal aperture. Moreover, vertical Kohler illumination requires a low power light source (2 watts LED is used) and illuminates only the imaging area of the leaf (~ 1 mm in diameter). Further, a vertical illumination limited to the imaging area of the leaf does not interfere with the stomata functions [8]. Importantly, it is possible to expose the upper epidermis of the leaf to the sunlight during stomata imaging. In the field, a leaf was fixed mechanically by a leaf holder and a leaf cap was used to restrict the ambient sunlight during imaging (Fig. 5D and 5D’).

### 3.4 Leaf Holder and Real-time Field Microscopy

For continuous monitoring of lower epidermis stomata, it is essential to hold the leaf surface parallel to the microscope-imaging plane. Leaf clip holders used in the measurement of leaf parameters such as chlorophyll contents (Heinz Walz GmbH, Germany) and electrical capacitance block the lower epidermis layer of a leaf [28]. Hence, these leaf clip holders are not suitable for continuous imaging of abaxial stomata. Therefore, a novel leaf holder has been designed and fabricated with biocompatible polymethylmethacrylate (acryl) and polylactic acid (PLA) material as shown in Fig. 2 and Fig. 6 [23–26].

**Fig. 6.**
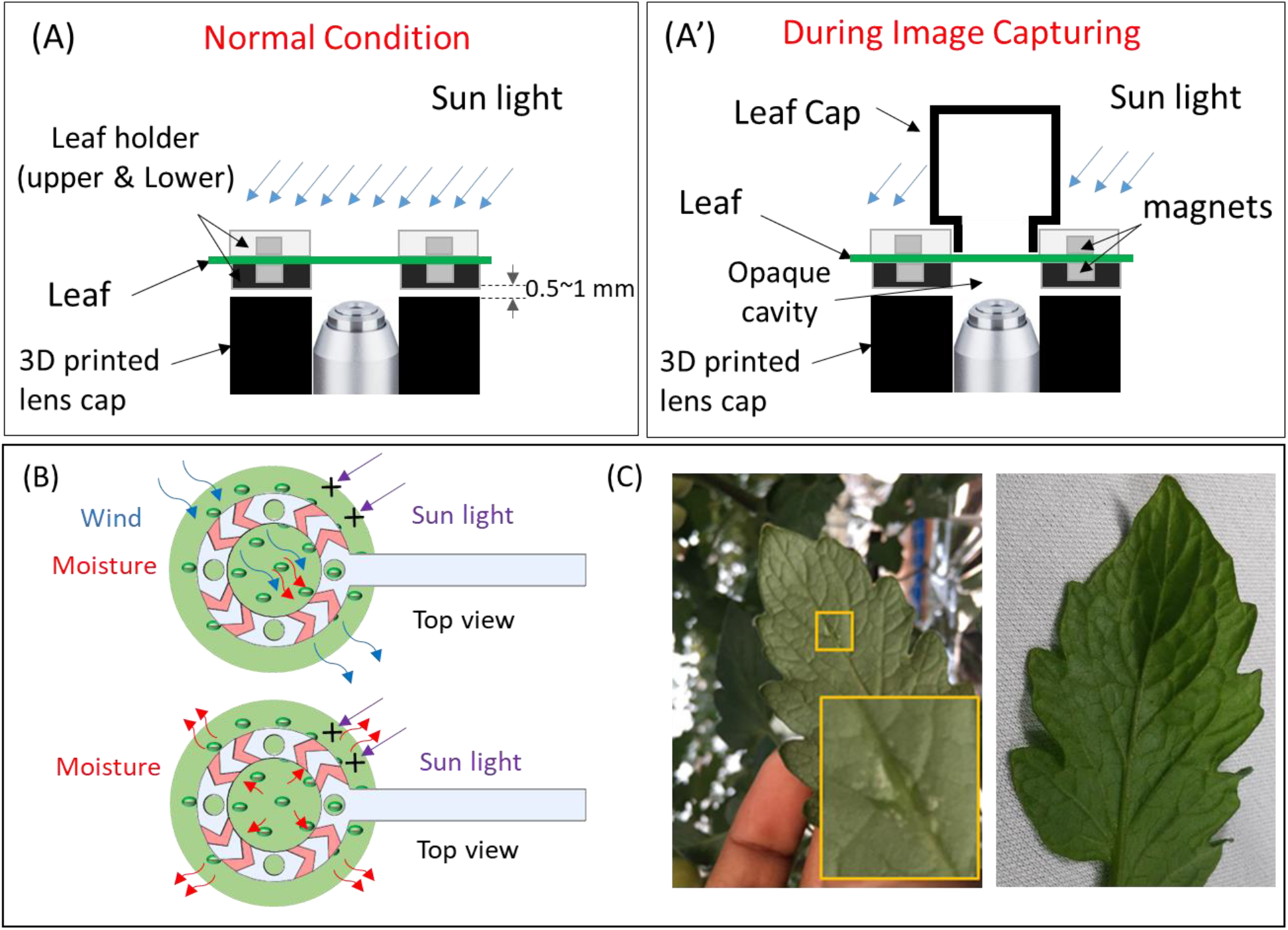
Leaf holder: a device to hold and stabilize the leaf for long-term stomata imaging. Schematic diagram of a leaf holder (A) in normal condition and (A’) during the imaging of stomata. (B) Schematic presentation of V-shaped notches allowing the wind blow and moisture across the leaf area under investigation and restricting the sunlight for the same area. (C) Water vapors accumulation in the night when lower leaf holder ‘2’ (left leaf) vs leaf holder ‘3’ (right leaf) was used. No physical damage was observed by leaf holder after continuous monitoring.

Our leaf holder allows the upper surface of a leaf open to the sunlight and the lower surface of the leaf to the wind simultaneously. Thick transparent acryl sheet of upper leaf holder holds the leaf in a position and allows the sunlight on the upper epidermis of a leaf, a predominant stimulus affecting the stomatal aperture [29]. A black color lower leaf holder restricts the ambient light reaching to the sample from backside [25, 26]. Further, to avoid the interference of ambient light reflected from lens surface with the optical system of the microscope, the lens was covered with a black color cap made of PLA material. However, the use of a lower leaf holder together with a lens cap forms an opaque cavity and restrict the wind to pass through the lower epidermis surface. In addition, stomatal water vapor is unable to diffuse into the environment. Combination of both events affect the stomatal conductance and result in a deposition of water droplets on lower epidermis.

Further, deposited water vapor affects the stomata functions as well as microscopy of stomata as shown in Fig. 6C (left leaf). Therefore, to allow the wind at abaxial side of the leaf, 2 mm wide V-shape notches were patterned on lower leaf holder. V-shape notches allow the wind and restrict the ambient sunlight pass through the abaxial side of a leaf. Small neodymium magnets embedded in upper and lower leaf holders, does not affect the sunlight intensity reaching the leaf epidermis. In addition, no visible physical damage on the leaf surfaces has been observed when the leaf was sandwiched between upper and lower lea holder for more than 24-hours (refer Fig. 6C (right). During microscopy, the upper surface of the leaf was closed by leaf cap to restrict the interference of ambient sunlight with the optical system of the microscope (Fig. 2C and Fig. 2C’). In such a way, the leaf was exposed to the ambient light throughout the experiment and high-quality images of stomata were obtained at regular interval by using lens cap without damaging the leaf.

### 3.5 Real-time Stoma Monitoring and Effect of Sunlight Intensity

Real-time observation of stomatal functions is relevant to understand the optimization of plant physiological processes over million years of evolution [3, 8, 30]. Changes in the stomatal aperture through external stimuli such as intensity of sunlight, temperature and humidity of surrounding environment and wind speed, is a result of the bidirectional flow of ions across the plasma and tonoplast membranes [31]. However, existing techniques are unable to quantify the changes in the stomatal aperture associated with environmental factors in a real-time manner under natural conditions.

To understand the effect of sunlight intensity on stomatal aperture, real-time monitoring of abaxial stoma was performed under sunlight in a field at the interval of 15 minutes. Prior to stomatal imaging, the leaf has been anchored by a customized leaf holder. Leaf holder attached with a X-Y-Z translation stage holds the leaf surface parallel to the imaging plane of the microscope and reduce the leaf movement in the windy environment throughout the experiment. Wind speed was negligible since the experiment was performed inside the greenhouse. A leaf, facing in the direction of sunrise, was selected to monitor the dynamics of stomatal aperture. However, in afternoon and evening, the leaf was under the shadow of other leaves and/or nearby tomato plants. Weather was clear and cloudy at the day of the experiment. Changes in the size of the stomatal opening (in μm^2^) in response to the variation in the sunlight intensity is shown in Fig. 7. It has been observed that the size of stoma opening is a function of the sunlight intensity falling on the leaf compared to the maximum sunlight intensity near about. The intensity of sunlight less than 5,000 lux triggers the opening of stomata and further, higher intensity of light actuates the opening of stomata. The presence of clouds (CL) blocking the sun affects the ambient sunlight intensity as well as the intensity of light close to the leaf (refer clouds (CL) 1, 2, 3 and 4 in Fig. 7B.). If the clouds block the sunlight for a long duration, such an event affects the stoma opening, (refer CL 2 in Fig. 7B). However, a latency between change in sunlight intensity and resulted change in the stomatal opening has been observed. If the leaf of the interest is in shadow during the clouds blocking the sun, the intensity of sunlight close to the leaf does not vary significantly, and stomata opening gets unaffected. Maximum stomata opening of 80.3 μm^2^ has been measured.

**Fig. 7.**
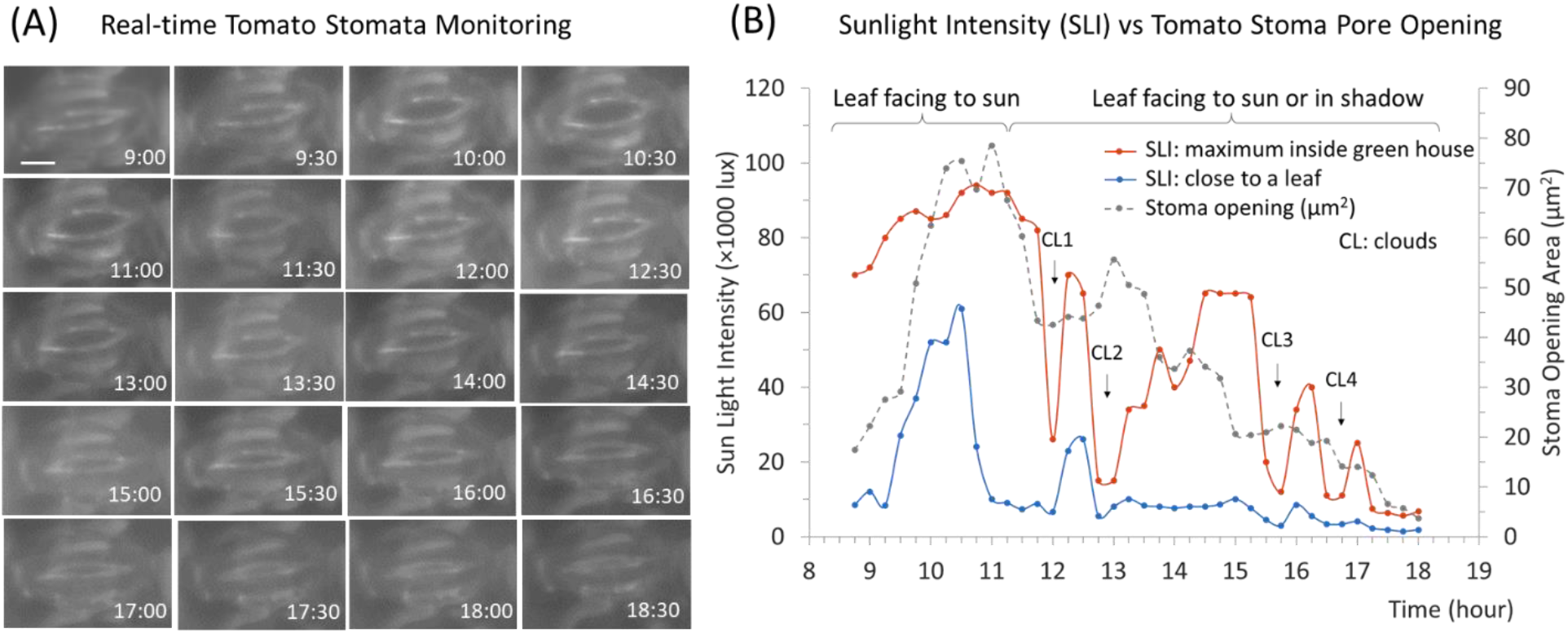
Real-time field monitoring of tomato stomata in sunlight (A) Imaging of stomatal opening at an interval of 15 minutes. Scale bar: 10 μm (B) Stomatal opening as a function of sunlight intensity. Stoma was responsive to the sunlight intensity falling on the upper epidermis compared to the maximum sunlight intensity of ambience inside the green house. The leaf under investigation was facing to sun in morning hours. However, during afternoon and evening it was under shadow of other leaves of the plant. The maximum stomatal opening of 80.3 μm^2^ has been observed after maximum sunlight intensity recorded parallel and close to the leaf surface.

### 3.6 Stomata Mapping of a Plant

Plant leaves coordinate stomatal opening/closure in response to light stress by sending and receiving rapid systemic signals [32]. Stomata mapping of four leaves exposed to the different intensities of sunlight at a time were conducted in six-week-old well-watered tomato plant as shown in Fig. 8. Leaves were detached from the plant at 1 pm and microscopic imaging of stomata was performed immediately. Leaf 1 and 2 were in shadow. However, leaf 3 and 4 were facing the sun partially and completely. The intensity of sunlight parallel and close to leaf surface was 2000 lux in case of leaf 1 & 2 and 13,000 lux & 60,000 lux for leaf 3 and 4 respectively. It has been observed that most of the stomata in leaf 1 and 2 were closed due to the low intensity of sunlight. However, for leaf 3 where the intensity of sunlight was 13,000 lux, stomata were partially open. Moreover, leaf 4, which has a maximum intensity of sunlight of 60,000 lux, stomata, were wide open. Results implied that, stomatal opening is a direct function of sunlight intensity falling on the leaf and dependent on the location of a leaf in respect to the light source including other factors such as wavelength of light source, stomatal density, leaf age, wind velocity etc. [5, 29, 33, 34]. Hence, it can be concluded that the contribution of a leaf for photosynthesis and transpiration depends on its location in respect to the sun especially for a plant having a high density of leaves. In addition, the plant porosity (total area of stomatal opening/ leaf area) changes continuously throughout of the day according to the geographical location of the plant in different weathers.

**Fig. 8.**
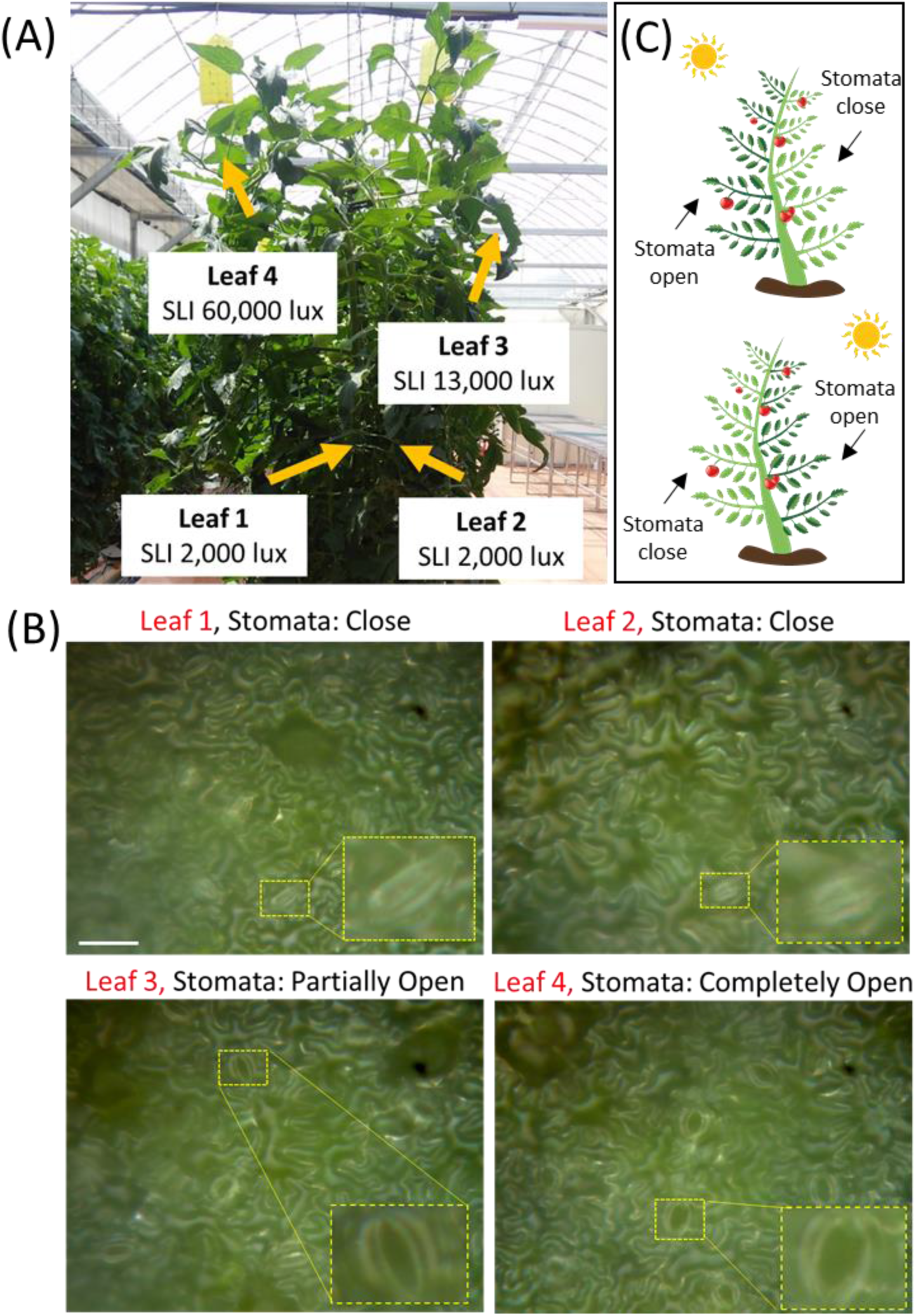
Stomata mapping of leaves. Leaves were detached from the plant and microscopy was performed immediately. (A) Location of Leaf 1, 2, 3 and 4 with respect to sun location. Leaf 1, 2, 3 and 4 were exposed to the sunlight intensity of values 2,000 lux, 2000 lux, 13,000 lux and 60,000 lux respectively. (B) Stomata were found to be close In case of leaf 1 and 2. Both leaves were in shadow and exposed to the low intensity of sunlight (2,000 lux). However, stomata in leaf 3 and 4 were either partially or completely open since both leaves were exposed to moderate and high sunlight intensity (13,000 lux and 60,000 lux). (C) A schematic of opening and closing status of stomata with respect to the position of sun. Scale bar: 50 μm

### 3.7 Quantification of Stomatal Density and Changes in Stomatal Geometrical Features in Response to the Sunlight Intensity

Static and dynamic stomatal properties control the stomatal conductance and determine the rate of transpiration and photosynthesis [35–37]. Real-time quantification of stomatal static property (stomatal density) and dynamic property (stomatal opening) in response to the sunlight intensity has been performed. An experiment was conducted on a terminal leaf of the fourth petiole of a six-week-old tomato plant. To distinguish the stomatal complex from other epidermis structures, stomatal counting was performed when stomata were widely opened. Stomatal density was 27±2.15 (n=5) in 350 μm × 262 μm FOV or 282 stomata per mm^2^.

Stomatal dynamic properties such as aspect ratio, circularity index and area of stomatal aperture were analyzed using multiple microscopic images in a given FOV (Fig. 9). Since, tomato leaf has a non-planar surface, the use of multiple microscopic images focused at different parts of the leaf in a given FOV provided an accurate result of stomatal dynamic properties. Status of lower epidermis stomata at 7:00 am and 11:30 am when the intensity of sunlight was 3,500 lux and 62,000 lux respectively are shown in Fig. 9A. When SLI on the leaf was low, stomata had a sub-micrometre opening. However, at high SLI, stomata were wide open. Observation of stomatal opening at regular intervals was conducted throughout the day and stomatal dynamic properties were quantified after measurement of stomatal opening as shown in Fig 9B.

**Fig. 9.**
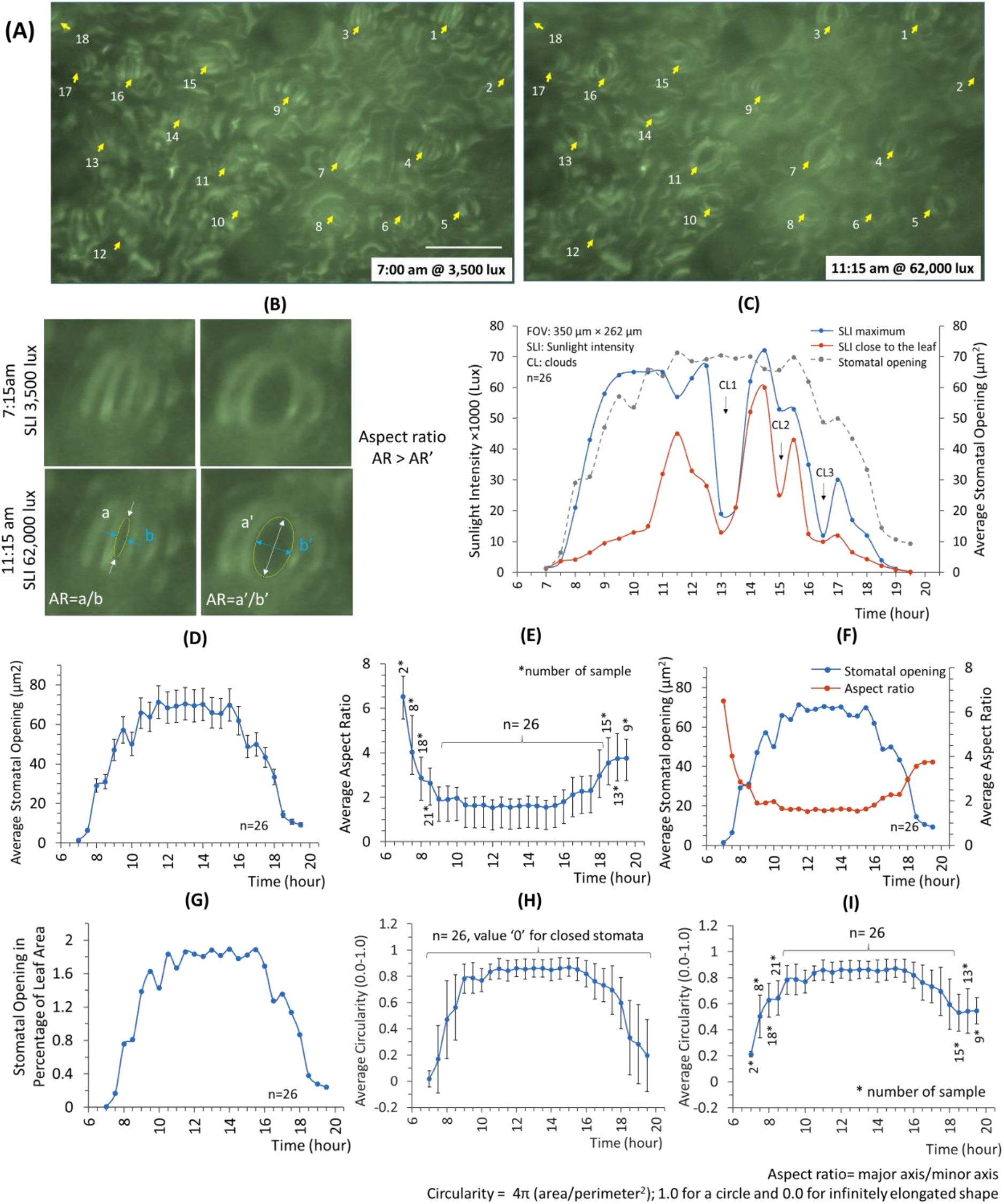
Quantification of stomatal density and stomatal opening geometrical features such as average/total stomatal opening area, aspect ratio and circularity of four-week-old tomato plant intact leaf under sunlight in a field. (A and A’) Status of the stomatal opening of an intact tomato leaf at 7:00 am and 11:15 am when the intensity of light close and parallel to the leaf surface was 3,500 lux and 62,000 lux respectively. Most of the stomata were close at 7 am, however, at 11:15 am; due to the high intensity of light all stomata were opened. To distinguish the stomatal complex from other structures of the epidermis, stomatal density was quantified when stomata were open. Scale bar 50 μm. (B) stoma opening status and measurement of geometrical features in microscopic images at 7:30 am (partially open) and 11:30 am (completely open) when SLI on the leaf was 4,600 lux and 45,000 lux respectively. (C) Time wise mapping of stomatal opening/closing as a function of SLI on a sunny/cloudy day. Stomatal opening reached close to the maximum when SLI on a leaf was more than ~15,000 lux. The average opening area remained to be similar through the day until SLI value on a leaf was more than ~15,000 lux. Time wise, average stomatal opening measurement. All stomata (n=26) in microscopic FOV (350 μm × 262 μm) were considered. For closed stomata, zero opening area was used for calculation of average stomatal area and standard deviation. (D) Average stomatal opening and total stomatal opening in the percentage of leaf area. Stomata start to open before the sunlight intensity reaches 10,000 lux. 20,000 lux SLI is sufficient to open the stomata completely. On average maximum stomatal opening area was ~75 μm^2^. However, the stomatal opening area may change according to the leaf and plant age. Total stomatal opening area was ~ 2% to the leaf area. (E) Average stomatal opening with standard deviation (F) Timewise measurement of average aspect ratio. Only open stomata were considered at the time of quantification. A declining and increasing trend in the value of aspect ratio was observed during opening and closing of stomata respectively. (G) Simultaneous representation of average stomata opening and aspect ratio. (H) Real-time monitoring of total stomatal opening in the percentage of leaf area. Circularity measurement of stomata (I) when all stomata (n=26) were considered for statistical analysis (zero value for closed stomata) (J) and only open stomata were considered.

Since, stomatal opening is a function of the SLI falling on the leaf instead of ambient SLI, average stomatal opening (n=26) did not reach its maximum value (~71 μm^2^) till 11:00 am even though ambient SLI (66,000 lux) was enough for maximum opening of stomata (Fig. 9C). A value of SLI on a leaf close to 15,000 lux was enough to have a wide opening of stomata. SLI higher than ~15,000 lux did not change stomata opening significantly although a fluctuation on stomatal opening has been observed at higher SLI on the leaf. However, latency has been observed in the change of stomatal opening in response to a change in SLI. Factors other than SLI such as type of plant, plant and leaf age, location of the leaf, ambient temperature, wind speed, atmospheric carbon dioxide etc., which also affect the stomatal opening, are not discussed here. CL1, CL2 and CL3 represent the clouds at different times, responsible for the reduction in maximum SLI inside greenhouse and SLI falling on the leaf. Further, an average aspect ratio of the stomatal opening (a ratio between the major axis and minor axis) together with the average stomatal opening was studied (Fig. 9B). It was observed that the aspect ratio of the stomatal opening decreases with increasing the stomatal opening area and wise versa (Fig. 9F). Only open stomata were considered for statistical analysis of aspect ratio since for closed stomata, aspect ratio will be a non-defined (0/0) mathematical expression. In addition to the aspect ratio, circularity index (CI, 4π (area/perimeter^2^), a measurement of roundness was also quantified. A maximum value 1.0 of CI represents the perfect circle; however, minimum value 0.0 represents the infinitely elongated object. During daytime shape of the stomatal opening was close to a circular shape however, in morning and evening when stomata start to open/close, CI index had a minimum value.

## 3. CONCLUSION

Here, we have demonstrated the direct real-time (RT) field imaging and monitoring of dynamics of the abaxial stomatal opening of a tomato leaf under natural conditions by a portable high-resolution reflected microscope without any physical or chemical manipulation of a leaf. Results establishes a direct relationship between the intensity of sunlight falling on the leaf and the area of the stomatal opening. Area of the stomatal opening increases with the increment in the sunlight intensity until maximum opening reaches and vice versa in well-watered plant. However, the age of leaf, location of the leaf, wind velocity, ambient temperature and other environmental factors may affect the concluded trend between stomatal opening area and intensity of sunlight at the leaf surface. In addition, stomatal density, changes in geometrical features of stomatal aperture (aspect ratio, circularity), changes in porosity of a leaf area as per intensity of sunlight has been quantified. Submicron meter spatial resolution (488 nm) along with the small depth of view (~ 1.3 μm) provided by our portable microscope enabled us to observe the stomatal pores of a non-planar leaf surface from it early opening until the maximum possible opening even in presence of high-density hair like structures. Use of novel leaf holder allowed us to perform long-term monitoring of stomata closely to its natural conditions. Further, a stomatal imaging of plant leaves exposing to the different intensity of sunlight at a time provided the mapping of plant porosity (total area of stomatal opening/ leaf area) which changes continuously throughout of the day according to the geographical location of the place in different weathers. We believe to be the first to demonstrate the real-time stomatal dynamics in intact plant leaf at the field. In future, field monitoring of stomata behaviour considering other environmental factors and conditions will further increase our understanding of the plant parameter(s) connected with stomata dynamics.

## Acknowledgements

This research was supported by the Rural Development Administration (PJ012100022016), Republic of Korea and the Bio-Mimetic Robot Research Center funded by the Defense Acquisition Program Administration and by the Agency for Defense Development (UD130070ID). The fabrication was performed at the Inter-university Semiconductor Research Center (ISRC) at Seoul National University.

